# A flexible method for estimating tip diversification rates across a range of speciation and extinction scenarios

**DOI:** 10.1101/2021.11.02.466977

**Authors:** Thais Vasconcelos, Brian C. O’Meara, Jeremy M. Beaulieu

## Abstract

Estimates of diversification rates at the tips of a phylogeny provide a flexible approach for correlation analyses with multiple traits and to map diversification rates in space, while also avoiding the uncertainty of deep time rate reconstructions. Available methods for tip rate estimation make different assumptions, and thus their accuracy usually depends on characteristics of the underlying model generating the tree. Here we introduce MiSSE, a trait-free, state-dependent speciation and extinction approach that can be used to estimate varying speciation, extinction, net-diversification, turnover rates, and extinction fraction at the tips of the tree. We compare the accuracy of tip rates inferred by MiSSE against similar methods and demonstrate that, due to certain characteristics of the model, the error is generally low across a broad range of speciation and extinction scenarios. MiSSE can be used alongside regular phylogenetic comparative methods in trait related diversification hypotheses, and we also describe a simple correction to avoid pseudoreplication from sister tips in analyses of independent contrasts. Finally, we demonstrate the capabilities of MiSSE, with a renewed focus on classic comparative methods, to examine correlation between plant height and turnover rates in eucalypts, a species-rich lineage of flowering plants.

## INTRODUCTION

Molecular phylogenies, when scaled in relation to time, are powerful sources of data to understand the diversification dynamics of organisms and have become crucial in multiple areas of ecology and evolution (Wiens & Donoghue, 2004). As the trees themselves become bigger, more robust, and increasingly comprehensive (e.g. Beaulieu & O’Meara, 2018; Smith & Brown, 2018), the statistical tools used to infer macroevolutionary patterns from them also become more biologically realistic (e.g. Beaulieu & O’Meara, 2016). Modelling the dynamics of lineage origination and extinction through time have allowed us to understand, for example, how shifts in pollination and dispersal strategies are connected to changes in diversification rates of angiosperms (e.g. Lagomarsino et al., 2016; Vasconcelos et al., 2019; Reginato et al., 2020) or the role of environmental instability in the diversification of several vertebrate groups (e.g. Harvey et al., 2020; Morales-Barbero et al., 2021).

Recently, however, a renewed wave of criticisms regarding these methods calls into question whether diversification rates from time-calibrated trees of extant-only organisms should even be estimated at all (Louca & Pennell, 2020). While it is true that phylogenies are often used to address problems beyond their capabilities (Losos, 2011; Cooper et al., 2016; Uyeda et al., 2018), there is still a considerable amount of information that extant-only phylogenies can provide about the diversification process (Helmstetter et al., 2021; Morlon et al., 2022). For instance, O’Meara and Beaulieu (2021) demonstrated that state-dependent speciation and extinction (SSE) models can identify different likelihoods for the generating parameters of trees with different topologies but identical lineage through time plots. However, they also discuss how the uncertainty around parameter estimates increases as one moves from tip to root in the tree, due to the decreasing amount of information in ancestral rate reconstructions (O’Meara and Beaulieu, 2021).

A solution, then, could be to continue to model diversification dynamics, but focus only on rates estimated near the present or at the tips of the tree, rather than deep in time. Tip rate estimates, also referred to as species-specific diversification rates (Maliet et al., 2019), have additional advantages in their flexibility when testing for correlations between multistate discrete and continuous traits (Harvey & Rabosky, 2018; Title & Rabosky, 2019) and for providing a straightforward way to map diversification rates in space (e.g. Sun et al., 2020; Suissa et al., 2021). Several methods for estimating tip diversification rates have been proposed in the last decade (e.g. Jetz et al., 2012; Rabosky 2014; Maliet et al., 2019) and they tend to provide different levels of accuracy for the estimates depending on the underlying model generating the tree (Title & Rabosky, 2019). There are also multiple views on how to use tip rate estimates in trait-based correlation analyses, depending on how the tip rates themselves were estimated (Freckleton et al., 2008; Rabosky and Huang, 2016; Maliet et al., 2019). Of course, the underlying processes generating empirical phylogenies are unknown, and therefore developing a flexible method that estimates accurate tip rates under a broad range of speciation and extinction scenarios is desirable.

Here we formally describe MiSSE (“Missing State Speciation and Extinction”), an extension of the SSE framework that provides accurate estimates of various metrics of diversification rates at the tips of a tree under various speciation and extinction scenarios. We also show how MiSSE estimates can be used alongside regular phylogenetic comparative methods after a simple correction for pseudoreplication of tip rates, contributing to its flexibility in tip correlation analyses. We compare the accuracy of MiSSE against other popular approaches for estimating tip diversification rates, and further demonstrate its capabilities with an empirical example that examines the correlation between turnover rates and plant height in eucalypts (Eucalypteae, Myrtaceae), a diverse lineage of flowering plants. Finally, we argue why focusing on tip estimates can be advantageous in testing complex hypotheses of diversification and discuss some caveats of this approach and possible ways forward for modeling diversification.

## MATERIALS AND METHODS

### The MiSSE model

State-dependent speciation and extinction models expand the birth-death process to account for speciation (*λ*), extinction (*μ)*, and trait evolution (*q*, the transition rates between character states), estimating parameters that maximize the likelihood of observing both the character states at the tips of the tree and the tree itself (Maddison et al., 2007). SSE models were initially developed to overcome three main perceived shortcomings in the field: (1) the need for greater flexibility in tests of key-innovation hypotheses, which, at the time, typically involved sister-clade comparisons that could only measure differences in net-diversification and assumed constant rates within the clades under comparison (Barraclough et al., 1998); (2) the need to incorporate differences in speciation and/or extinction rates associated with a particular character state in analyses of trait and geographical range evolution (Goldberg et al., 2010; Goldberg & Igic, 2012; Ree & Sanmartin, 2018); and (3) the potentially confounding effects of different transition and diversification rates when looking at diversification or character evolution, respectively (Maddison, 2006).

All discrete SSE models exist within the following generalized ordinary differential equations (corresponding to eq. 1a-b in FitzJohn, 2012):

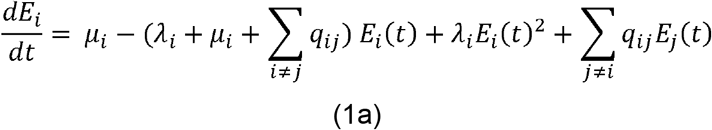

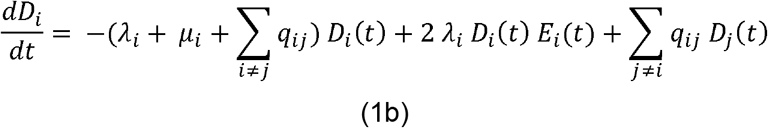

The probability E_*i*_ (*t*) is the probability that a lineage starting at time *t* in state *i* leaves no descendants at the present day (*t* = 0), and D_*i*_ (*t*) is the probability of a lineage in state *i* at time *t* before the present (*t* > 0) evolved the exact branching structure as observed.

With these ordinary differential equations, any number of states can be included in an SSE model. For example, in character-based models, such as BiSSE (“Binary-State Speciation and Extinction”; Maddison et al., 2007), *i* and *j* represent two observed states of a focal character (e.g. observed states 0 and 1). A potential issue in this case is that when comparing a simple model where there is no variation in rates among states against a model of trait-dependent diversification there is almost always strong support for a trait-dependent diversification process in empirical settings (Maddison & Fitzjohn, 2014; Rabosky & Goldberg, 2015). The HiSSE model (“Hidden-State Speciation and Extinction”; Beaulieu & O’Meara, 2016) partially corrects this issue by harnessing the properties of hidden Markov models to allow rate heterogeneity to depend not only on the focal trait, but also to be correlated with other factors that were not explicitly scored as character observations at the tips (see also Caetano et al., 2018; Nakov et al., 2019; Boyko & Beaulieu, 2021, 2022). Thus, with HiSSE, *i* and *j* represent the different observed and hidden states combinations specified in the model (e.g. observed states 0 and 1, hidden states *A* and *B*).

Although hidden state SSE models made hypothesis testing in diversification studies more realistic, existing SSE models are still generally used to understand the correlation of a particular observed trait or geographical range distribution and the diversification dynamics of a group. However, in reality, it is not a single factor but, rather, a combination of circumstances that are responsible for the heterogeneous diversification rates among clades in a phylogenetic tree (Donoghue & Sanderson, 2015; Nürk et al., 2020). To understand heterogeneous processes that arise from the effect of multiple traits on the dynamics of speciation, extinction, and trait evolution is one of the main utilities of hidden Markov models in phylogenetic comparative methods (Caetano et al., 2018).

It is natural, then, to completely drop the observed trait from the analysis and focus only on the impact of the “unobserved” traits, or the hidden states, in the diversification dynamics of a clade. This is what our MiSSE model, an extension of the HiSSE framework, is intended to do (see also Barido-Sottani et al. 2020 for a similar SSE extension in a Bayesian framework). MiSSE is a direct extension of HiSSE with the main difference between the two models being that with HiSSE, we have an observed character with states 0 or 1, as in BiSSE (Fig.1a), *and* hidden states *A* and *B* (Fig. 1b). Essentially, MiSSE operates in the same way, but simply ignores the observed states altogether and performs the calculations of the hidden rate classes directly (Fig.1c).

**Figure 1:**
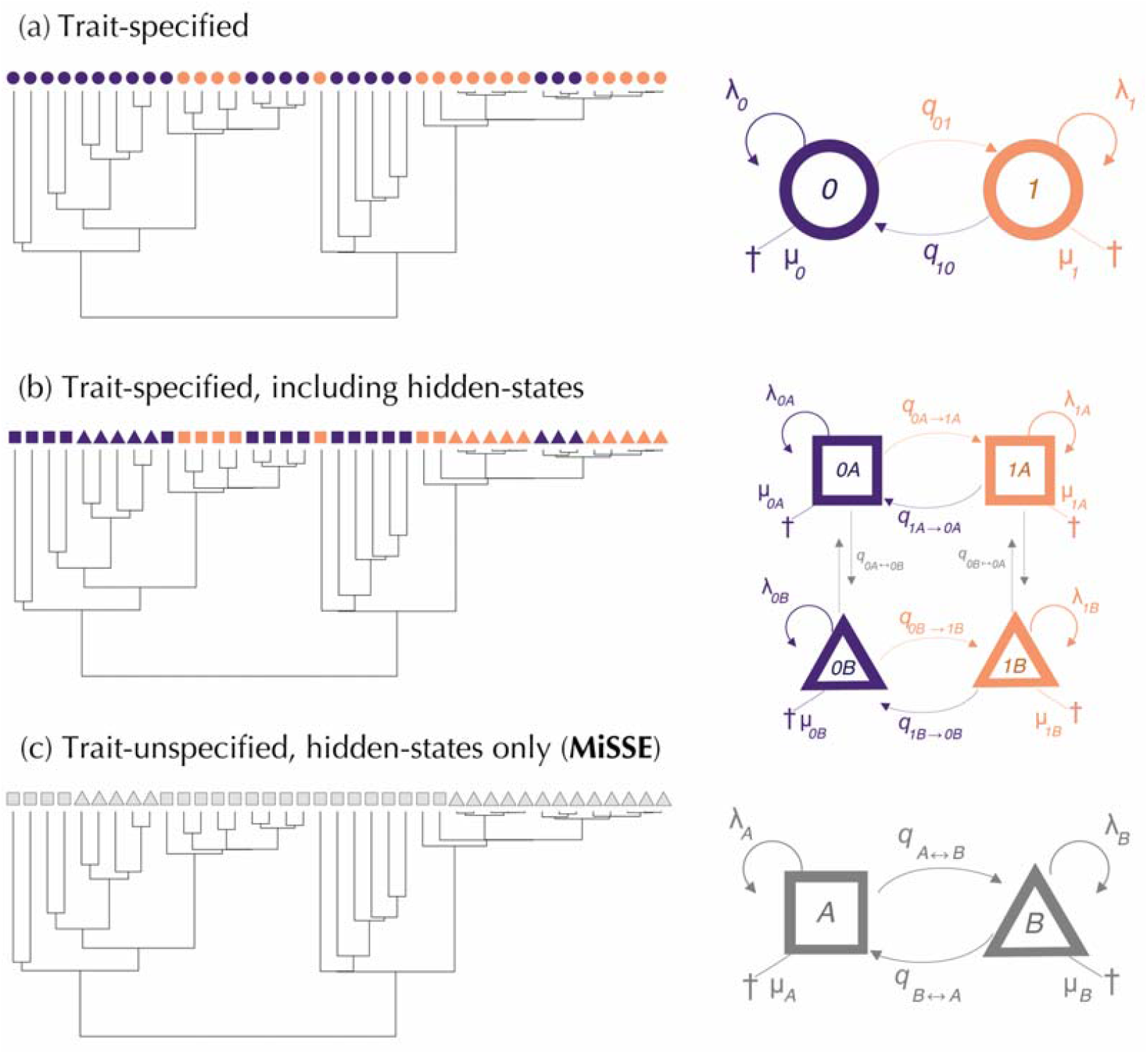
Diagrammatic representations of state-dependent speciation and extinction models that: (a) use only observed trait data in parameter estimation (e.g. BiSSE), (b) use both observed trait data and hidden-states in parameter estimation (e.g. HiSSE, where observed and hidden are coded as numbers and letters and are combined to yield a four-state system); and (c) use only hidden states in parameter estimation (e.g. MiSSE).

The assignment of particular hidden state to each tip is based on probabilities of the data, which, in the case of MiSSE, is just the structure of the tree. This process is like the maximum likelihood calculations in other hidden state methods. These calculations are described thoroughly in Caetano et al. (2018), so here we provide only a brief overview. The first step is to search for parameter values that maximize the probability of observing the tree at the root by marginalizing over all possible assignment of rate classes at internal nodes and along branches. This search consists of a bounded stochastic simulated annealing algorithm, run for 5000 iterations, followed by a bounded subplex routine that runs until the maximum likelihood is found. Alternating between a stochastic optimization routine, followed by a “greedy” hill climbing routine, helps ease MiSSE away from finding local optima.

Suppose that, in an iteration, rate class *A* has λ = 0.1 and μ = 0.05, and rate class *B* has λ = 0.2 and μ = 0.1. The node subtending the branch subtending a tip could have started in *A* giving rise to an *A* or *B*, or it could have started in *B* giving rise to an *A* or a *B*. Each of these scenarios has a probability associated with them. The sister edge has the same set of scenarios and its own set of probabilities. These probabilities are combined at the nodes (which includes the speciation rate to account for the speciation event at a node) and carried down the tree in the same way as in other SSE models. Once at the root, marginal probabilities for whether the root is in *A* or *B* based on a set of rates can be calculated, as well as the overall log-likelihood that these rates produced the observed tree.

At this point, all other branches, nodes, and tips are assumed to be in all possible states. The second step, then, is to take the MLE of the rates and determine which of the states at a given node and given tip is more likely than any of the other possible states. For that, MiSSE simply chooses a node (or tip), fixes it to be in rate class *A*, traverse the tree, and calculates the overall likelihood. Then it does the same for that same node, but this time it fixes it to be in rate class *B*. At the end, MiSSE calculates the marginal probabilities by dividing the probability that a node was in each state divided by the sum of the probabilities across all states. In that way, every tip and node have a probability of being in both states *A* and *B*, but often one state will be more probable than the other. For example, in a clade where speciation occurred more rapidly, the higher λ associated with *B* will be a better fit to the shape of that part of the tree, and so the marginal probabilities will reflect higher support for *B*. In another clade, where speciation is slower, lower λ will be the better fit, so the marginal probabilities will reflect the higher support for hidden state *A* in that part of the tree. And in other clades they may be uncertain, given frequent transitions in and out each hidden state.

Note that *A* and *B* are arbitrary labels that can and will shift positions at the tips in different runs. What really matters is the parameter combination that underlies each label in each run, which should not change their MLE between runs. Note too that MiSSE is still a model of tree and character states like other SSE models, and so the topology matters to assign tips to the correct (hidden) states. Because it uses data from the topology as well as branch lengths, MiSSE likelihood can distinguish between trees that have the same lineage through time plot, but different topologies (O’Meara & Beaulieu, 2021). Also note that, contrary to previous SSE models, transition rates (q) among rate classes are always set equal. They are informed in the output, but they have no direct interpretable biological meaning like the transition rates in BiSSE, which represent the frequency of state changes among observed character states, or in GeoSSE, which can be interpreted as the frequency of dispersal out of a biogeographic area. Our current implementation will continue to treat these rates as fixed until it is clearer whether transition rates are practically feasible in MiSSE. An implementation where transition rates are allowed to vary would be ideal given that shifts between rate classes are likely to occur at different speeds throughout the evolution of a group. Differential diversification rates have a major effect on the tree (since this affects exponential growth of clades) while differential transition rates are likely (but not guaranteed) to have a less substantial effect, so for now we use the data only to estimate differences in the former.

MiSSE is available within the R package *hisse* (Beaulieu & O’Meara, 2016). Some details of MiSSE’s implementation differ from other SSE models also implemented in *hisse*. These differences are summarized at SM1 (Supplementary Material) and readers are also encouraged to follow the example code available as SM2 when using MiSSE for their empirical analyses.

### Comparing the accuracy of tip rate metrics across different speciation and extinction scenarios

We compare the accuracy of tip diversification rates estimated by popular methods using a set of simulated trees extracted from Title and Rabosky (2019) and Maliet et al. (2019). Title and Rabosky (2019) used 5200 trees compiled from previous literature and simulated under eight different models as a benchmark set to test the accuracy of different tip rate metrics across a range of speciation and extinction scenarios. For speed, we selected a random sample of 10% of their original set of simulations, i.e. 521 trees, for our analyses. We then excluded trees simulated under the “multi-regime, constant-rate birth–death” of Meyer and Wiens (2018) because those were represented by only two trees in our set. From Maliet et al. (2019), we extracted their 40 trees simulated under the ClaDS2 scenario for the comparisons, the same model available in the recent data-augmentation implementation of the ClaDS model (Maliet and Morlon 2022). We then excluded five trees where the median height of branch lengths was above a threshold of one trillion units of time because they could be out of the numerical limits for some methods (e.g. BAMM’s MCMC chains struggled to reach convergence in those cases). Finally, seven additional trees were excluded due to modelling limitations in one of the methods (see details in SM3; Supplementary Material). The final dataset comprises 547 trees of different sizes, though most of them are < 500 tips (Table 1).

**Table 1:**
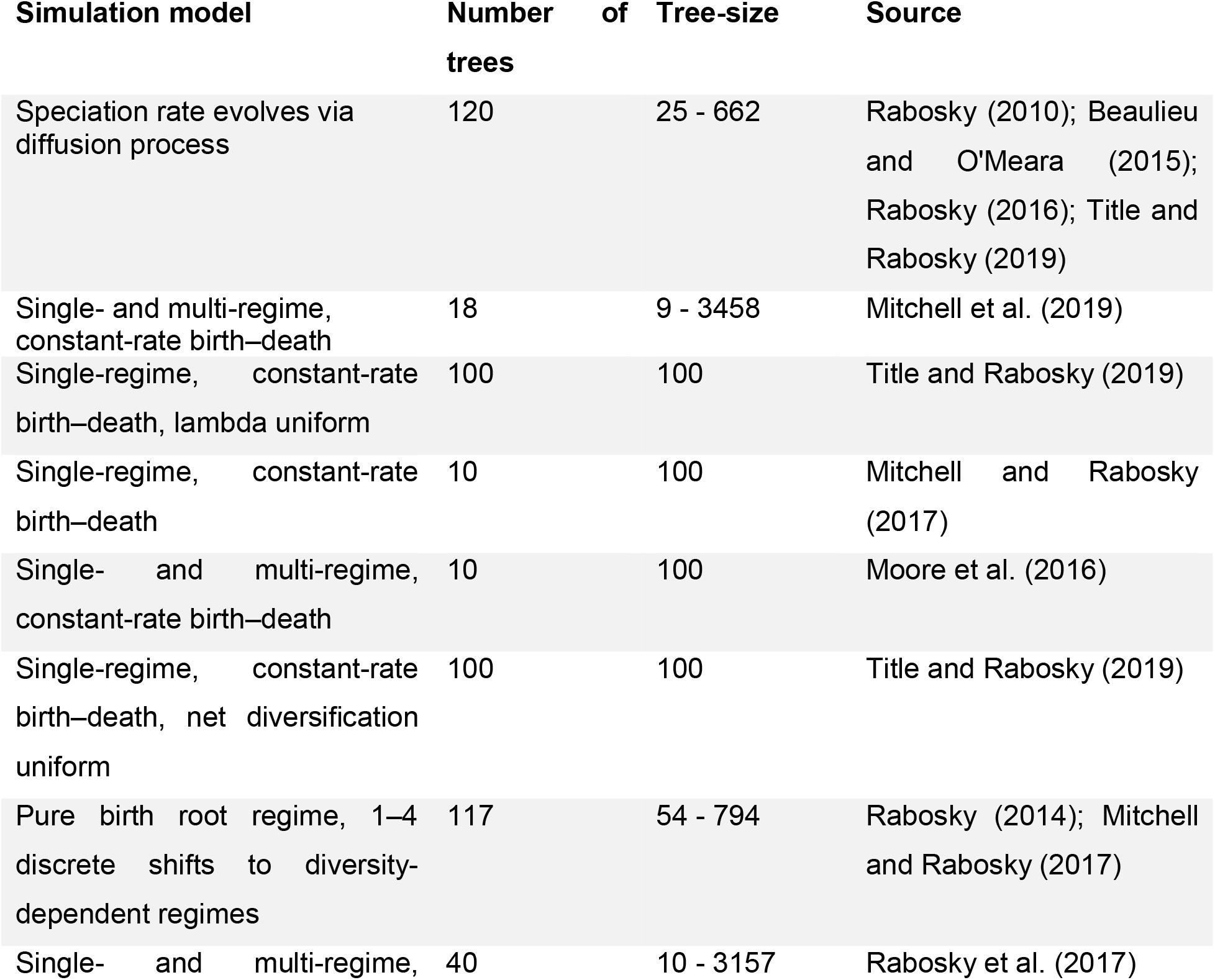
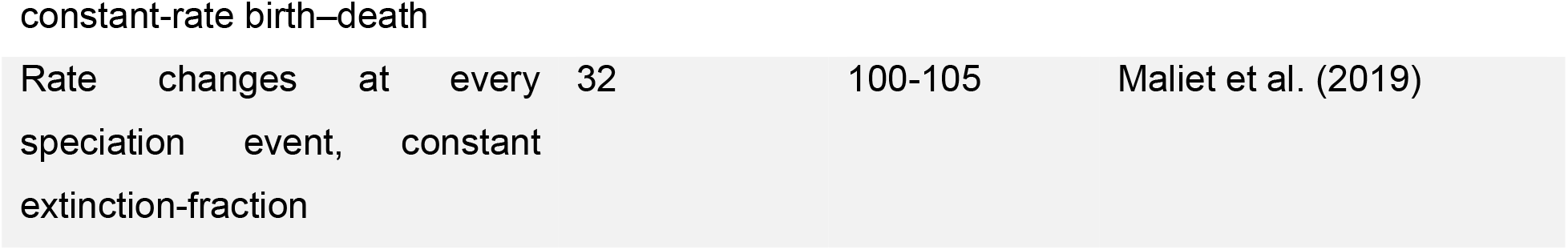
Simulated trees used in comparisons between tip rate metrics.

Six different tip-rate metrics were compared: DR statistics (Redding & Mooers, 2006), BAMM (Rabosky, 2014), node density (ND; Freckleton et al., 2008), the inverse of terminal branch length (TB; Steel & Mooers, 2010), ClaDS (Maliet et al. 2019; Maliet and Morlon 2022), and our MiSSE model. For MiSSE, we summarized tip rates in the following ways: (1) tip-rates estimated from the model with overall lowest AICc (i.e. the “best” model; MiSSE_best_); and (2) tip-rates estimated by averaging all models according to their Akaike weight (MiSSE_average_) (see SM3, Supplementary Material, for details on the settings for other methods).

We followed the same statistical metric as Title and Rabosky (2019), namely, we compared mean absolute error, given by the formula 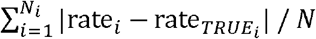, and the RMSE is given by 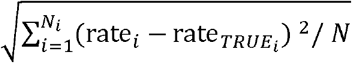 where *i* is a single tip and N is the total number of tips in all trees of a given type of simulation. In all cases, the lower the error values the more accurate is the metric. We assessed the accuracy of five parameters: speciation (λ), extinction (μ), and the orthogonal transformations of these, namely, net-diversification (r = λ - μ), turnover (τ = λ + μ) and extinction fraction (ε = μ / λ). Note that TB, ND and DR are non-parametric (i.e. not based on an underlying model) and only estimate λ. We then used the function posthoc.kruskal.conover.test from the R package PMCMR (Pohlert, 2014) to perform a pairwise test for multiple comparisons of mean rank sums and calculate whether errors are significantly different between metrics, assuming a significance value of p < 0.05.

### Empirical example: plant height and turnover in Eucalypts

We demonstrate the capabilities of MiSSE estimates for analyses of tip correlation with an empirical example. Body size is considered an important trait in studies of animal evolution (Cooper & Purvis, 2010) and it has been interpreted as a potential correlate of diversification rates in vertebrates (Cope’s rule; FitzJohn, 2010). A similar argument can be made for plants (Boucher et al., 2017). Through a complex link between rates of molecular evolution, fecundity, and population size, life span and generation time are expected to be negatively correlated with the number of speciation and extinction events on a per time basis (Stebbins, 1974; Petit & Hampe, 2006); i.e. slower turnover rates. Although these correlations are expected in theory, surprisingly few studies have compared them in practice using model-based approaches (e.g. Boucher et al., 2020). The MiSSE framework allows us to easily test this correlation by using regular comparative phylogenetic methods and tip-rates as a response variable.

We examined turnover rates in relation to plant height, a proxy for lifespan (Westoby, 1998), in eucalypts, one of the most distinctive components of the Australian landscape (Wilson et al., 2005). Eucalypts are members of the tribe Eucalypteae in the flowering plant family Myrtaceae, a group that includes some of the tallest trees in all angiosperms (e.g. *Eucalyptus regnans* can reach up to 120 m height) (Wilson et al., 2005; Nevill et al., 2010). Eucalypts are also unusual for being considered a relatively speciose (c. 800 species) lineage of trees. It is thought that the tree habit tends to decrease speciation rates, which is offered as reason that clades comprised of predominantly tree growth forms are frequently found to be less diverse than their herbaceous or shrubby relatives (Petit & Hampe, 2006). However, many eucalypts species are also large shrubs to treelets between 1.5 and 5m height (EUCLID, 2015), which led to the hypothesis that the radiation is driven primarily by the smaller representatives of the clade (Petit & Hampe, 2006). Here we test this idea by contrasting tip turnover rates with plant height in this diverse clade of angiosperms.

We used the “ML1” time calibrated phylogenetic tree from Thornhill et al. (2019), which covers 716 out of the c. 800 species of eucalypts. We then collected data on plant height for 673 species available at EUCLID Eucalypts of Australia Edition 4 (2015) (EUCLID, 2015). All measurements represent plant maximum height and are given in meters. Again, we use the default implementation of MiSSE, that is, stop.deltaAICc=10 and chunk.size=10, to estimate tip turnover rates in Eucalypteae, pruning redundant models with the function PruneRedundantModels. Next, we estimated tip turnover rates averaging all models according to their AICc weights with the function GetModelAveRates.

To test correlations between plant height and tip turnover rates, we used the function TipCorrelation, which performs a regression-through-the-origin of positivized phylogenetic independent contrasts (PICs, Felsenstein, 1985) between tip rates and a continuous trait. This function also gives the option of pruning out PICs from two tips that are sister to each other and share the same branch length to the direct ancestral node (i.e. “cherries”), before regressions. We reasoned that because these sister tips theoretically inherit the exact same rate class probabilities in MiSSE they may: (1) present identical tip rates, forcing the slope of the regression to be close to 0 since their PICs will be 0; and therefore (2) constitute pseudoreplicates in the analyses. Note that, in this approach, we prune PICs and not individual sister tips. Pruning tips might not be adequate because it would affect all other PIC calculations in the tree; it also can make the results sensitive to which tip is pruned and gives less information for computing contrasts deeper in the tree. All functions mentioned above are available in the R package *hisse* (Beaulieu and O’Meara, 2016). Rates and maximum plant height were log scaled before analyses so that they conformed with Brownian motion evolution (Felsenstein, 1985; Garland et al., 1992).

## RESULTS

### Simulation studies

Comparisons between tip rate metrics using simulated data show that, in general, MiSSE estimates accurate tip rates across a range of speciation and extinction scenarios (Fig. 2). We present the results for mean absolute error, though RMSE results are practically the same (SM4; Supplementary Material). Our results show that TB is significantly less accurate than other methods in all simulation scenarios, and, in eight out of the nine simulation scenarios, there was no significant difference in error measurements between ND and DR. In all simulated scenarios, results show that the model-based approaches MiSSE, BAMM and ClaDS tended to be significantly (p < 0.01) more accurate than the non-parametric methods TB, ND, and DR for speciation rates, the only parameter estimated by the latter three metrics (see SM5; Supplementary Material for all pairwise comparisons).

**Figure 2:**
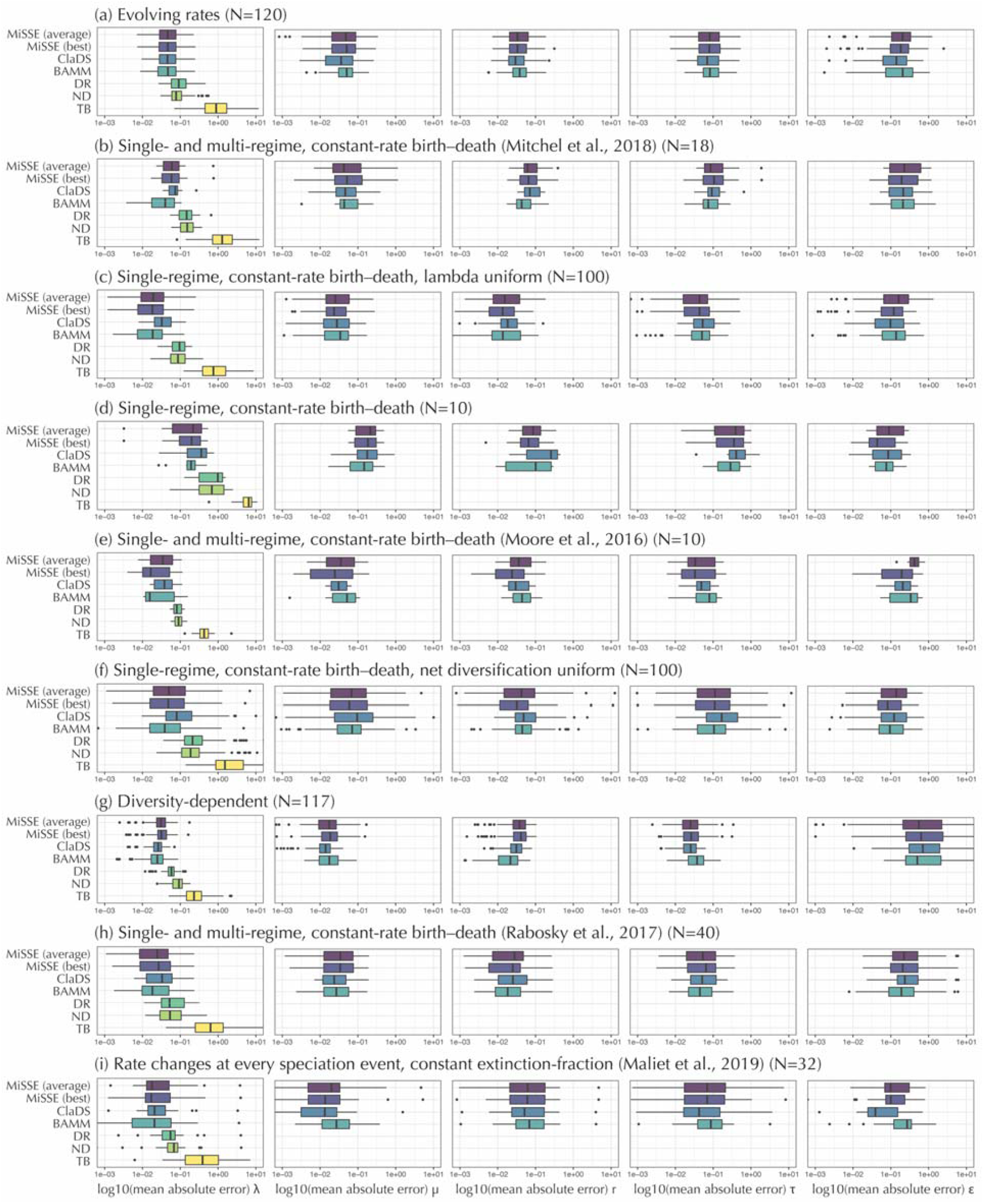
Boxplots showing the distribution of mean absolute error between true and estimated tip rates using five previously described metrics and our MiSSE model. Each data point corresponds to the mean absolute error between all tips in one tree. λ = speciation rate, μ = extinction rate, = turnover rate, and r = net-diversification rate, = extinction fraction. Letters between brackets highlight significantly different error measurements.

Accuracy among the three model-based approaches for the five diversification parameters estimated at the tips varies depending on the model used to generate the simulated tree. Significant differences in accuracy are observed for extinction fraction in the “speciation rate evolves via diffusion process” scenario (Fig. 2a), where ClaDS is significantly more accurate (p = 0.04) than MiSSE_average_ and BAMM, but not than MiSSE_best_ (p = 0.32). Significant differences in accuracy between the estimates of the MiSSE_best_ model and the AICc-weighted average of the MiSSE_average_ model were not observed in any other simulated scenario; therefore, the two results are henceforward treated as a single MiSSE category. In both the “single-regime, constant-rate birth–death, lambda uniform” (Fig. 2c) and the “single-regime, constant-rate birth–death, net diversification uniform” (Fig. 2f) scenarios, ClaDS estimates for speciation at the tips are significantly less accurate (p < 0.01) than the other model-based approaches. ClaDS estimates also perform significantly poorly than MiSSE (p < 0.01 and p = 0.049 when compared to MiSSE_best_) for estimates of net-diversification and turnover rates in those scenarios. BAMM and ClaDS are significantly more accurate than MiSSE for speciation in the “Pure birth root regime, 1–4 discrete shifts to diversity-dependent regimes” scenario (p < 0.01; Fig. 2g). In the same scenario, ClaDS also tends to be more accurate than other methods for extinction rates (p < 0.01) and BAMM for net-diversification (p < 0.01), but BAMM is also significantly less accurate for turnover rates (p < 0.01) than the other methods in this scenario. ClaDS is significantly more accurate than BAMM for estimates of extinction fraction in trees simulated under the model “rate changes at every speciation event, constant extinction-fraction” (p < 0.01; Fig. 2i), but does not significantly differ from MiSSE for any parameter in that scenario.

### Tip-correlation between plant height and turnover

Our empirical example using tip rate correlations with MiSSE show that, despite the idea that smaller plants have driven the radiation in eucalypts, maximum plant height appears uncorrelated with turnover across the tips of the tree (R^2^ < 0.001, Fig. 3c). The general lack of correlation is mainly because: (1) higher turnover rates are not always restricted to clades of small shrubs and treelets, as it can be seeing by comparing the distribution of tip-rates and plant height (Fig. 3a,b); and (2) plant height (Fig. 3b) appears to be a much more labile trait than turnover rates, regardless of the methods used to infer tip-rates (Fig. 3a,b). We note that pruning PICs from sister tips from the tree minimally changes the slope of the regression line in this example and it does not change our main conclusion that turnover and plant height are likely uncorrelated in eucalypts, given that contrasts of maximum plant height explain less than 0.1% of the contrasts of turnover. In any event, the inclusion of PICs from sister tips can force the slope of the PIC regression towards 0 in other tip rate comparisons and therefore they should be pruned before analyses.

**Figure 3:**
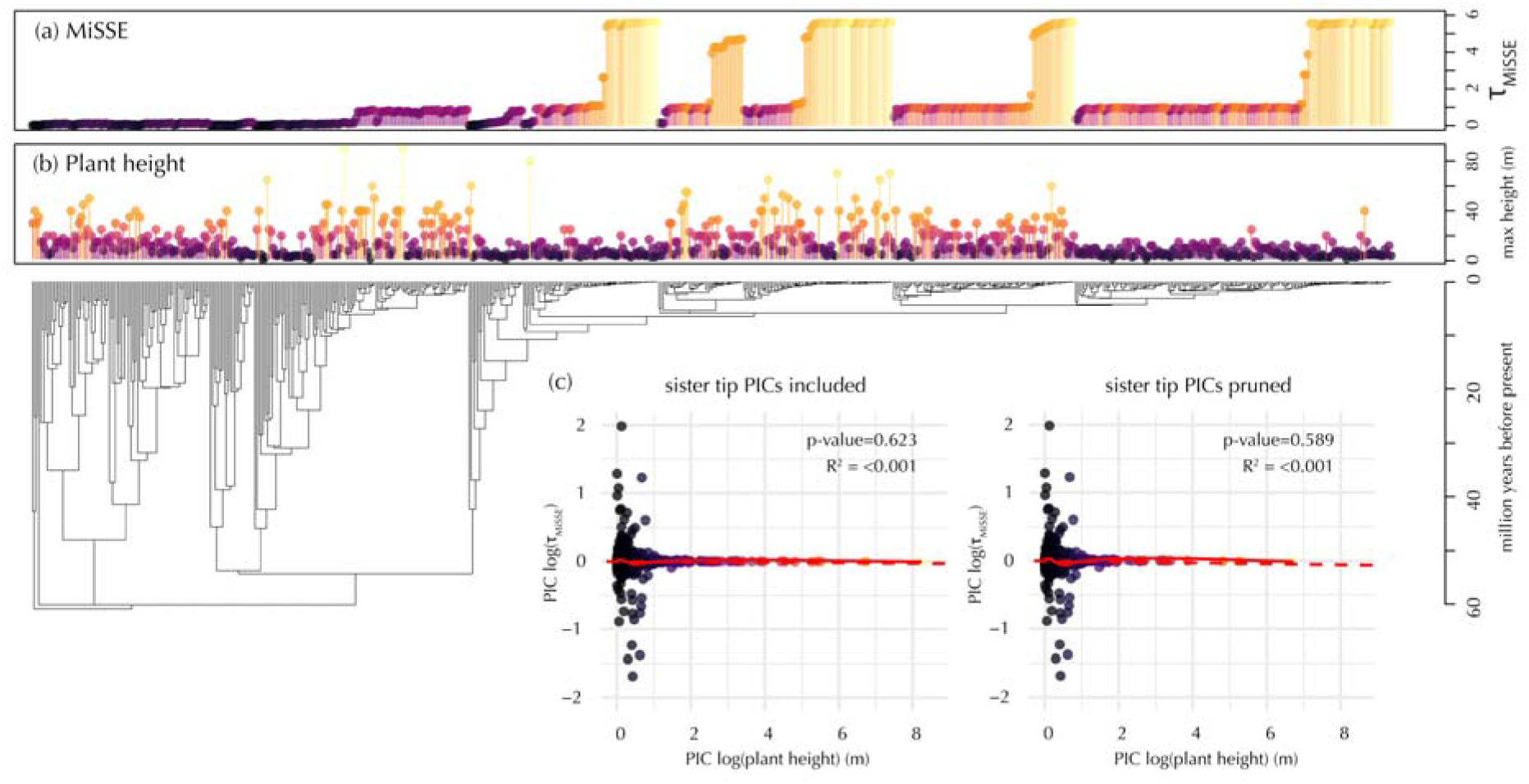
Comparison between maximum plant height and tip turnover rates estimated by MiSSE in Eucalypteae. (a) MiSSE turnover rates at the tips; (b) Maximum plant height in meters; warmer colors indicate higher values in both cases. (c) Regression through the origin between PICs of MiSSE turnover rates and plant height, with PICs from sister tips included and pruned. Solid line: smoothed line using the loess method; dashed line: regression line.

## DISCUSSION

### MiSSE as a flexible model-based approach to estimate tip diversification rates

Here, we describe our “trait-free” MiSSE framework and show how it can be used to estimate accurate diversification rates at the tips of the tree. Our results show that the difference in accuracy in parameters estimated by averaging all MiSSE models or using only the best MiSSE model seem to be minimal and may depend on the tree (Fig. 2). However, as discussed in previous publications (Beaulieu & O’Meara, 2016; Caetano et al., 2018), the user is strongly encouraged to focus mainly on parameter estimation rather than which model fits “best” when using MiSSE, since the former is more informative about the biology of a clade. We also note that, although extinction fraction and turnover are sometimes synonymous in the literature (see SM6, Supplementary Material, for a short terminology survey), we use turnover rate as an explicit measure of events per units of time, and a parameter that is explicitly orthogonal to extinction fraction. We emphasize this difference because many diversification hypotheses may be better described by turnover rates rather than net-diversification or extinction fraction (e.g. Vrba, 1993; Vasconcelos et al., 2022; our empirical example). Also, given that extinction and speciation should tend to correlate through time (Marshall, 2017), net-diversification should tend to approach 0. In those scenarios, and as long as extinction is different from 0, turnover can naturally become a better metric for understanding diversification dynamics nuances than its isolated components. We note, however, that when extinction rate is equal to 0, turnover rate collapses into speciation rate (i.e. there is no turnover). We also note that our concept of turnover is different from the one used in the island biogeography literature (e.g. Simberloff 1974).

Key differences between MiSSE and analogous methods are highlighted by our analyses of simulated data. Although estimates from the three model-based approaches, MiSSE, BAMM and ClaDS, tend to provide generally accurate estimates of diversification parameters at the tips, the three methods make very different assumptions. BAMM is based on inferences of discrete rate shifts, so that all tips after a shift inherit a similar rate (Rabosky et al. 2014). ClaDS, on the other hand, assumes that changes in diversification dynamics occur at every speciation event, so that the estimates at the tips are allowed to be highly heterogeneous in relation to one another (Maliet et al. 2019). MiSSE may also have some shift-like properties, as in BAMM, in areas where the probabilities of being in different rate classes change abruptly. However, because at every moment in the tree, including at the tips, there is a combined probability of being in all possible rate classes, the evolution of diversification rates becomes smoother, allowing it to pick up some rate heterogeneity at the tips, as in ClaDS. Closely related tips do have similar rates because it is likely that they will be in the same rate class, but they do not inherit the absolute same rate.

A visual inspection of individual results (Fig.4; all plots in SM7, Supplementary Material) show that differences in accuracy observed in our comparisons probably result from these operational differences between model-based approach. For instance, because BAMM works with discrete shifts, this method tends to be more accurate when the true model assumes that few large discrete shifts of diversification rates occurred in the tree. In those scenarios, models like MiSSE and ClaDS may perform comparatively worse because they may infer rate heterogeneity where there is none (Fig. 4d). On the other hand, BAMM can underestimate rate heterogeneity when the true model produces highly heterogeneous tip rates in the tree. In those cases, even if the mean error is low, this method may fail to capture rate variation in small clades with particularly high or particularly low rates (Fig.4a,c). Similarly, ClaDS may overestimate variation in scenarios where rate heterogeneity is nonexistent, resulting in higher error (Fig,4b). Because MiSSE works as an intermediate between BAMM and ClaDS, we argue that it can become a model-based approach that is more flexible in identifying areas of rate heterogeneity or homogeneity at the tips of the tree. Importantly, even in scenarios where MiSSE is less accurate, it tends to capture areas with faster and lower rates in the tree correctly. In “diversity dependent” scenarios, for example, MiSSE may be less accurate because it can overestimate speciation rates of clades with higher speciation rates, since it does not explicitly allow speciation rates to slow down along a given terminal branch (as in BAMM). However, it tends to be similar to other methods in identifying where variation in rates occurs (Fig. 4d).

**Figure 4.:**
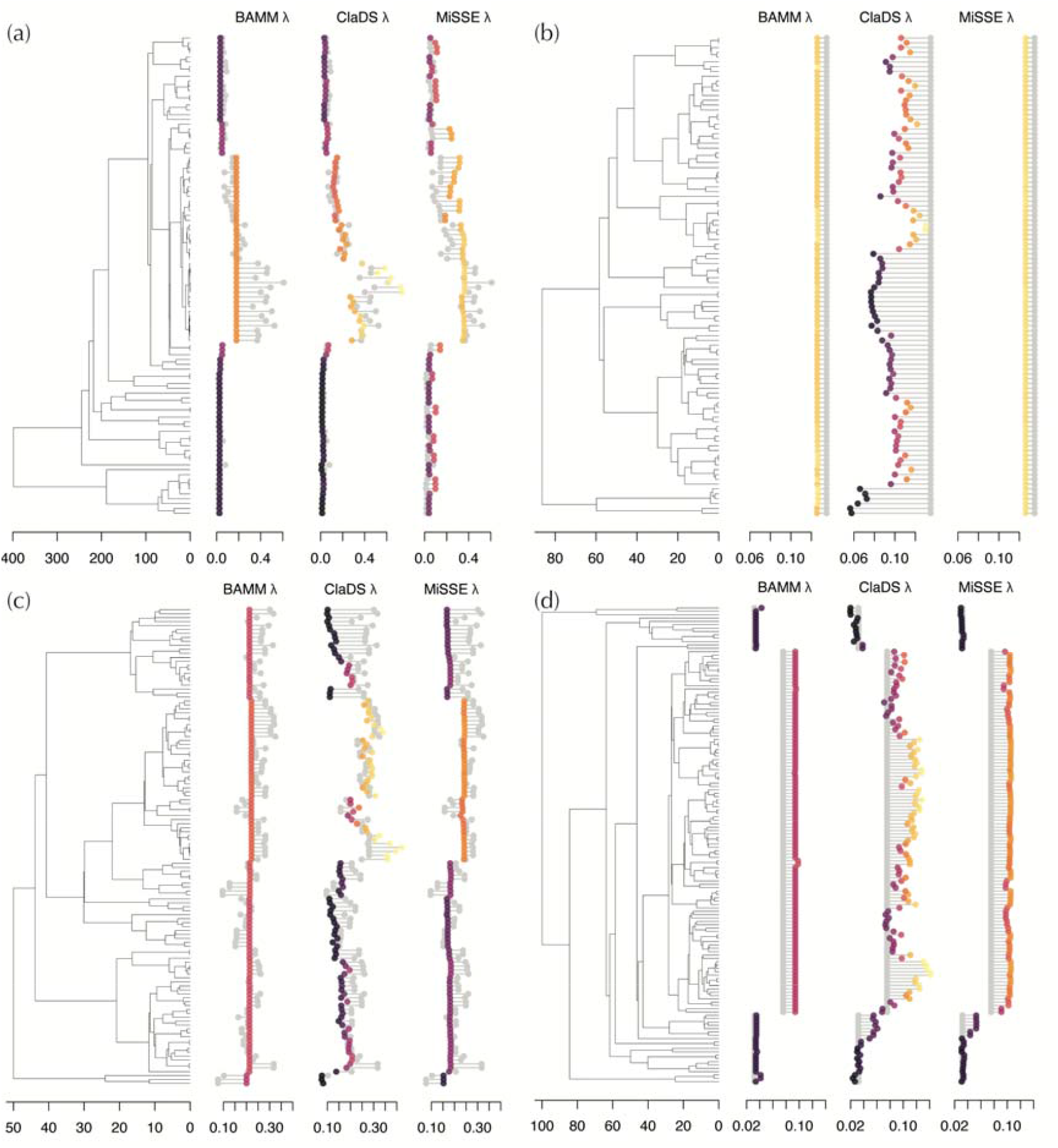
A comparison between rates estimated using different model-based approaches (colored dots, warmer colors indicate higher values) and true rates (gray dots) at the tips of the tree in four different simulated scenarios: (a) “rate changes at every speciation event”; (b) “speciation rate evolves via diffusion process”; (c) “single-regime, constant-rate birth–death, lambda uniform”; (d) “pure birth root regime, 1–4 discrete shifts to diversity-dependent regimes”.

### Plant height and turnover rates are uncorrelated in eucalypts

Our empirical analysis demonstrated that the correlation between plant height and turnover rates is weak within Eucalypteae. The use of plant height as a proxy for longevity is common in plants, but it may be that plant height is not the best proxy for longevity in eucalypts. Tall eucalypt trees are often native to productive environments with fertile soils (Pryor, 1976; Thornhill et al., 2019) and it may be that they present fast life cycles for their size, as observed in other parts of the world where soils are also fertile (Russo et al., 2008). In fact, many eucalypt trees are known to be fast-growing plants (e.g. Barnard et al., 2003; Almeida et al., 2004) and fast growth is linked to higher mortality through the growth-survival trade-off (Reich, 2014), which could potentially lead to higher turnover rates even in lineages of tall trees (Baker et al., 2014). If that is the case, perhaps plant height is not the trait that best captures the variation that we sought to explore in our hypothesis. Other proxies, such as leaf mass area and seed size (Westoby, 1998; Wright et al., 2004) may be a better proxy of longevity in these scenarios and should also be tested.

Alternatively, if plant height is, in fact, a good proxy for longevity in eucalypts, it may be that longevity is generally uncorrelated with turnover in the clade. A similar trend of uncorrelated body size and speciation rates has been observed in other groups (FitzJohn, 2010; Boucher et al., 2020), but it is difficult to ascertain if the lack of correlation is a particularity of the groups that have been analyzed so far or a general rule. Given the heterogeneity of evolution, identifying what is a rule and what is an exception in macroevolution requires analyses of multiple natural replicates. Similar tests in several clades would be ideal to rule out this possibility. A different approach would be to use QuASSE (FitzJohn, 2010) to construct a model of turnover and height; the disadvantage of this is the need to model the relationship and the (current) impossibility of including additional observed or unobserved factors.

One benefit of MiSSE (and similar methods) is that, even though our hypothesis was not supported, we still have new information about the tips: turnover rate (and the other diversification parameters if one wishes). In the same way that looking at the correlation of two traits may generate new hypotheses to test (Boyko and Beaulieu, 2021), having estimates of turnover rate at tips as in Figure 3 may also lead to new ideas about the factors leading to higher turnover rates in some eucalypts.

### Best practices in tip rate estimation

MiSSE is intended to be used to estimate diversification rates at the tips of the tree. Tip rates are, technically, an emergent property of how diversification dynamics are modeled along the branches of the tree (Freckleton et al., 2008; Title & Rabosky, 2019). However, rate estimates are not unique per tip species, so species should not be analyzed in isolation. In other words, a tip with extinction-fraction of 0.9, for instance, does not necessarily indicate that that tip is about to go extinct, but rather that a lot of closely related species are relatively short-lived in comparison to the rest of the tree. Tip rates can then be seen as a “snapshot” of the diversification potential of the species in a clade. In the MiSSE framework, this diversification potential may be interpreted as the combined effect of the many (hidden) states evolving in the tree, which are represented by the different rate classes.

From a mathematical standpoint, there is an advantage in using tip rates because it is at the tips where the certainty around the probability of being in a particular rate class is the highest (O’Meara & Beaulieu, 2021). The certainty around these probabilities decreases as one moves towards the root of the tree, and so the ability to point out changes in rate classes also decreases. Of course, the model *does* use information from the whole phylogenetic tree in the calculations of tip-rates, but as one moves towards the present in the tree, the more information is available to reconstruct those rates and thus we are more likely to be certain about the probabilities of being in different rate categories (see O’Meara & Beaulieu, 2021).

The intuitive question that follows is how far back in the past one can go to interpret how the observable diversification potential at the tips came to be and how far into the future can we use it to make predictions about how clades will diversify. The answer is, again, not straightforward and it may depend on the size and age of the phylogenetic tree under analysis. Tip rates will not be able to tell us about mass extinction events deep in the past as these are arguably only measurable from the fossil record (Barnosky et al., 2011), but they may represent a good reflection of diversification dynamics above the species level in times slices close to the present. We therefore recommend great caution with literal interpretations of rate classes deep in time. That is why although MiSSE can “paint” the averaged rate reconstruction along the branches of the tree (i.e. with the function plot.misse.states in the R package *hisse*), inferences of rates through time should not be interpreted literally. The “painting” should be used just to visualize the rates inferred with MiSSE and should not be used to support narratives of past diversification dynamics for a group. Interpreting past rates is risky given limited information. Similarly, modern drivers of diversification dynamics (climate change, invasive species, habitat destruction, and more), which affect especially extinction rates at the present, will not be reflected in the rate estimates returned by MiSSE or similar methods.

Another issue is the non-independence of the tip rates. The identical rates for sister tips have been discussed above, but even non-sister tips may have somewhat correlated rates. It is important to realize that though a phylogenetic model was used in the estimation of tip rates, they themselves are not corrected for phylogeny (which is a reason we used independent contrasts above to compare turnover and plant height). There is also limited information: a resolved tree of 100 taxa has 198 edges. Though MiSSE, BAMM, ClaDS, and other methods give 100 tip estimates, there is not enough information to take each one as known with great certainty, nor to draw a strong conclusion based on a rate shared by a few taxa.

### Caveats and persistent issues in modeling diversification dynamics

Tip-diversification rates are advantageous for their flexibility in trait related diversification analyses and for avoiding the uncertainty of rate reconstructions in deep time. However, there are several caveats that users should consider when using these methods, including MiSSE: (1) Clade-specific sampling fraction was found to lead to an incorrect likelihood behavior in *any* diversification method despite its appeal (Beaulieu, 2020). Contrary to other similar methods, MiSSE deliberately does not include an implementation for clade-specific sampling fraction, so all sampling fractions are global and only one sampling fraction is given for the whole tree. Imputation using stochastic polytomy resolvers can work as an alternative solution when dealing with incomplete phylogenies (e.g. Chang et al. 2020; but see Rabosky 2015) (2) How species are defined is the most relevant when looking at tip rates. For diversification rate models, the data come from the distribution of branching events across the phylogeny. Since most of these events are nearer the present, lumping and splitting taxonomic entities at the species level will have a greater impact on tip-rate estimation. We suspect that the methods discussed herein are particularly sensitive to taxonomic subjectivity. (3) There may be still issues related to ascertainment bias and underestimation of extinction rates (Beaulieu & O’Meara, 2018). In that sense, the larger and broader taxonomic sample one uses to test diversification hypotheses, the more biologically realistic extinction rates will tend to be. Ideally, one would be able to correct the extinction estimates in smaller trees, but that is still not possible with the current implementation of MiSSE nor in any other model we are aware of. So, even though extinction rates are estimated accurately relative to the simulated data, they may still be biologically unrealistic in small and young trees. (4) Finally, given that these models do not account for mass-extinction events, it is unclear what impact such events will have on tip-rate estimation in any approach currently available.

## CONCLUSIONS

Time-calibrated phylogenetic trees have been important sources of data to understand fundamental aspects of the evolution of organisms. MiSSE provides a novel tool to explore this, and its flexibility in estimating accurate tip rates across a broad range of speciation and extinction scenarios makes MiSSE a powerful alternative for diversification analyses with correct likelihoods. There remain many caveats and cautions about its use, but it is an additional tool to understand diversification processes, including a focus on parameter of perhaps great biological relevance such as turnover rate. We suggest that tip-rates estimated by MiSSE will be useful to several questions that were previously addressed by other SSE models. Its versatility is appealing to explore integrative questions linking traits, geographical range distribution, or both in time slices close to the present.

## Author contributions

T.V., B.C.O. and J.M.B. designed the study. J.M.B. led software development, T.V. and B.C.O. led the simulation study and T.V. led the empirical example. T.V. wrote the first draft and all authors contributed to the final writing of the manuscript.

## Acknowledgements

The authors thank members of the Beaulieu lab for discussion. The authors are also grateful to D. Rabosky and two anonymous reviewers for comments that greatly improved an earlier version of this manuscript. This work was funded by the National Science Foundation (grants DEB–1916558 and DEB–1916539). The authors have no conflict of interest to declare.

## Data Accessibility Statement

All code and data required to reproduce the analyses in this manuscript can be found at https://github.com/bomeara/missecomparison and in the Dryad folder linked to this publication: []

